# Chromosome-level genome assembly of *Protandrena* (*Anthemurgus*) *passiflorae* (Hymenoptera: Andrenidae), a host-plant specialist bee

**DOI:** 10.1101/2025.02.05.636715

**Authors:** Katherine A. Parys, Rena M. Schweizer, Ligia Benavides, Scott Geib, Sheina Sim, Jay D. Evans, Michael G. Branstetter

## Abstract

The passion flower bee, *Protandrena* (*Anthemurgus*) *passiflorae* (Robertson) is a monolectic, host-plant specialist of the passionflower plant *Passiflora lutea* L. Using a single adult male individual, we generated long-read PacBio HiFi, HiC, and short-read RNA sequencing data to build a well-annotated, chromosome-level genome assembly for this species. The final nuclear genome is 249 Mb with 150x coverage and with most of the genome scaffolding into 12 chromosomes. The scaffold N50 is 21.4 Mb and the genome has a Benchmarking Universal Single-Copy Ortholog (BUSCO) score of 97.2% for 5991 hymenopteran genes. BRAKER3 annotation of the genome identified 12,098 genes and 15,353 total transcripts and found that 20.27% of the genome is made up of repetitive elements. We resolved a mitochondrial genome of 12.7 kb. The *P. passiflorae* genome represents one of only a few published andrenid bee genomes and one of the first monolectic bees. This new high-quality genome will serve as a valuable resource for investigating the genomic basis of specialization and for providing a useful resource for studying pollinator health and conservation.

## Introduction

Sequencing genomes for every species on Earth has become a major objective of biodiversity science in the 21st century (e.g., Rhie et al. 2021; Hotaling et al. 2021). Similar to the collection, preservation, and curation of specimens in natural history museums, capturing genomic information of diverse species and specimens helps us conserve and understand biological diversity on the planet. Once acquired, reference genomes can be used for a multitude of basic and applied purposes, including evolutionary and functional genomics, conservation genetics, breeding, and systematics. A recent initiative, called Beenome100, aims to generate reference genomes for diverse wild bees in the United States, focusing on species of agricultural, biological, and conservation importance. The project aims to provide genetic resources that will help protect bee diversity and the critical pollination service that they provide and to facilitate future research. This study presents a new genome for this effort.

The passion flower bee, *Protandrena* (*Anthemurgus*) *passiflorae* (Robertson, 1902), is a monolectic bee described as a small, black, robust and reniform bee that is approximately 7.5 – 8.5mm in length (Michener et al. 1994; Neff and Rozen 1995). Originally described as a monotypic genus, *Anthemurgus* is currently considered a subgenus of *Protandrena* (Bossert et al. 2022). This species is considered “rare” across North America in Michener et al. (1994) but can be locally abundant in areas where *Passiflora lutea* L., its host plant, is abundant, particularly in Mississippi (Parys et al. 2018). This species is known from central Texas, Arkansas, Kansas, Illinois and east to North Carolina in the United States (Figure 1D; Michener et al. 1994; Neff and Rozen 1995; Holland and Lanza 2008; Mitchell 1960) and was recently collected in Mississippi in 2018 (Parys et al. 2018). Neff and Rozen (1995) described detailed foraging behavior, nests, and larval characteristics, but the bee has received little subsequent study.

**Figure 1.**
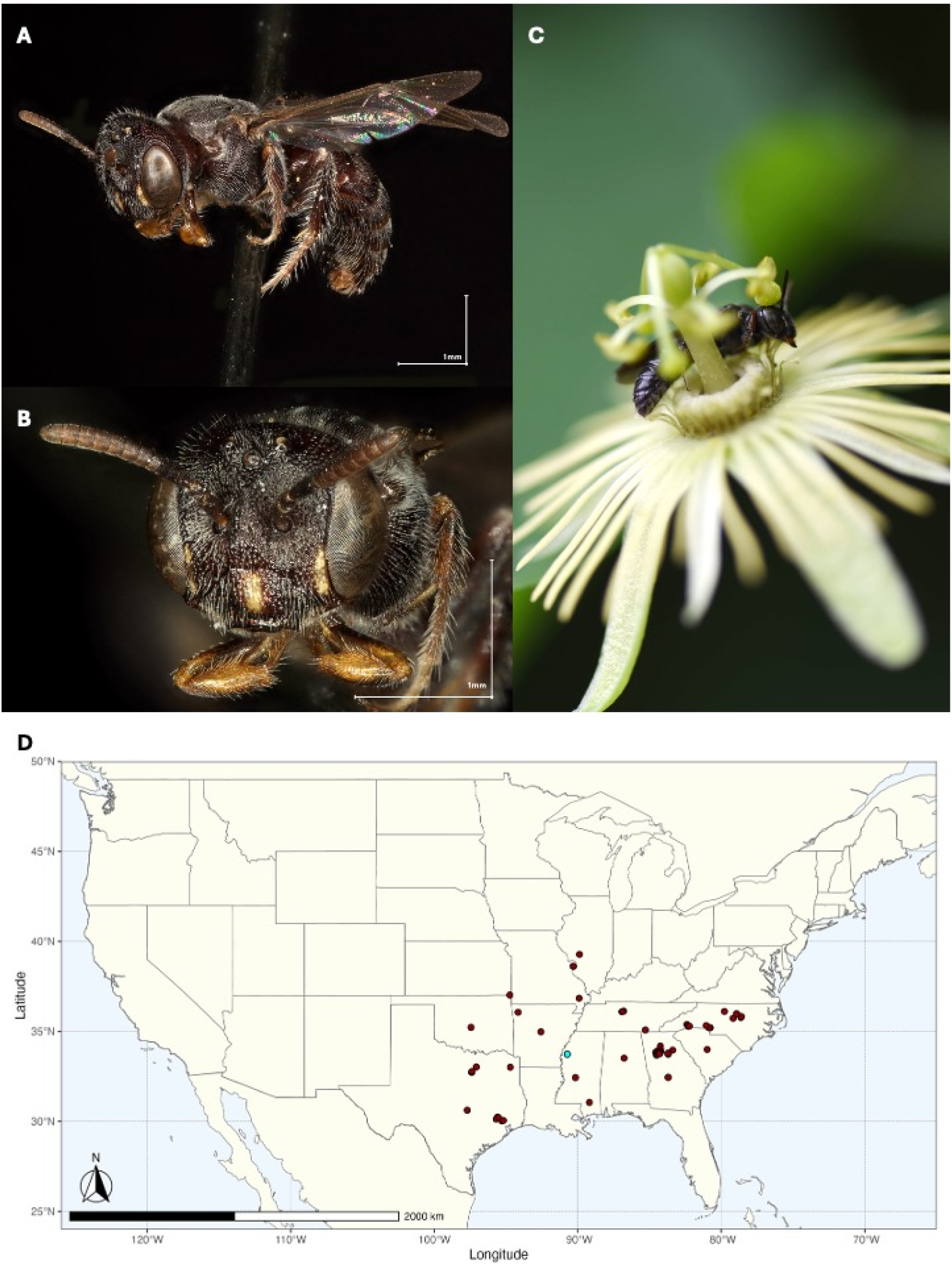
Lateral (A) and Head (B) images of voucher specimen of *Protandrena passiflorae*, deposited at PIRU. *Protandrena passiflorae* on its host plant, *Passiflora lutea* (C). Map of *P. passiflorae* distribution in the United States.

*Protandrena passiflorae* is one of the few truly monolectic bee species, known to solely forage on a single host plant species (Cane 2020). The yellow passionflower, *Passiflora lutea* L., is endemic to the southeastern U.S. and occurs in mesic open woodlands and vernal shaded areas from central Texas to Florida, and north to southern Kansas and Pennsylvania (Goldman and MacDougal 2015c). It is also a host plant for several species of *Heliconius* butterflies (Lepidoptera: Heliconiinae) across the southernmost parts of the range (Cardoso 2008; Estrada and Gilbert 2010).

Specialist bees are often thought to be more efficient pollinators than generalist visitors, but female *P. passiflorae* foraging for pollen are thought to act as pollen thieves on *P. lutea*, rarely ever contacting the stigmas during the process (Neff and Rozen 1995; Parker 1981). Host plant specialists are common in several groups of bees, but *P. passiflorae* is the only known bee to be oligolectic on a species in the family Passifloraceae (Rasmussen et al. 2020; Neff 2003). The pollen collection behavior of *P. passiflorae* appears to be unique to *P. lutea*, with no foraging observed on sympatric species with similar phenologies, *Passiflora incarnata* L. and *Passiflora affinis* Engelm. (Neff and Rozen 1995; Goldman and MacDougal 2015b, 2015a).

Here we present a high-quality reference genome for *P. passiflorae*, the first genome sequence for the genus *Protandrena* and subgenus *Anthemurgus* and one of only a few available genomes outside of the nominal genus *Andrena*. This genome is important because of the taxonomic position of the bee and its unique biology as a specialist pollinator of passion flowers. This genome will facilitate understanding the genetic basis of oligolecty/monolecty within bees, help elucidate andrenid phylogeny and evolution, and contribute to conservation of an endemic plant. Insect genomic initiatives like the i5k consortium, Ag100Pest, AgriVectors, and now Beenome100 (https://beenome100.org) are highlighting the benefits of creating high-quality and improved genomic resources and building open access tools to make them more accessible (Robinson et al. 2011; i5K Consortium 2013; Giraldo-Calderon et al. 2015; Childers et al. 2021; Saha et al. 2021; Isaacs-Thomas 2022).

## Methods and Materials

### Sample collection

Individuals of *P. passiflorae* were collected by net directly off blooms of *Passiflora lutea* in Cleveland, Mississippi, located in Bolivar County (Figure 1; 33.73521, -90.73244). The collection of five live males occurred on the 21st of July 2019, and bees were placed individually into cryo tubes, placed on ice, returned to the laboratory, and placed directly into a -80C freezer. We have designated specimen SIMRU20858 as the official specimen voucher and it has been deposited into the National Pollinating Insects Collection (NPIC) at the USDA-ARS Pollinating Insects Research Unit in Logan, UT (Figures 1A-C; unique voucher# SIMRU36530).

The distribution map (Figure 1D) was generated using R version 4.3.1 (R Core Team, 2023) and the following R packages: cowplot v. 1.1.3 (Wilke, 2024), ggrepel v. 0.9.5 (Slowikowski, 2024), ggspatial v. 1.1.9 (Dunnington, 2023), googleway v. 2.7.8 (Cooley, 2023), maps v. 3.4.2 (Becker et al., 2023), rnaturalearth v. 1.0.1 (Massicotte & South, 2023), rnaturalearthdata v. 1.0.0 (South et al., 2024), sf v. 1.0.16 (Pebesma, 2018; Pebesma & Bivand, 2023), tidyverse v. 2.0.0 (Wickham et al., 2019), with available distribution data (GBIF, 2024). Images of the voucher specimen (Figures 1A and 1B) were taken using a Keyence VHX-7000.

### DNA extraction and sequencing

DNA from a single adult male specimen (specimen# SIMRU36530) was used for all genome sequencing. For long-read sequencing, high molecular-weight (HMW) DNA was extracted from part of the abdomen using a Qiagen MagAttract HMW DNA kit (Qiagen, Hilden, Germany) and DNA quality was assessed using a Denovix DS-11 fluorometer/spectrophotometer (Denovix Inc, Wilmington, DE) for purity and quantification and a Fragment Analyzer (Agilent, Santa Clara, CA) for size assessment. The HMW DNA was sheared to 15-20 kb using a Megaruptor 3 (Diagenode, Denville, NJ, USA) and a single SMRTbell library was prepared using the SMRTbell 3.0. library preparation kit following manufacturer’s protocols for low input samples (Pacific Biosciences, Menlo Park, CA). The final library was sized with Ampure PB beads (Beckman Coulter, Brea, CA) to remove fragments less than 3 kb in length. The entire library was quantified using Qubit HS DNA reagents, sized on an Agilent Fragment Analyzer to determine molar concentration and then bound and loaded for sequencing on a single cell on a Pacific Biosciences Sequel IIe system. A HiC library was prepared from abdomen tissue. The proximity-ligated sequencing library was prepared from crosslinked tissue following standard protocols (adapted from e.g. Lafontaine et al 2021) using restriction enzymes DdeI and DpnII and subjected to PE 150 bp sequencing on an Element Biosciences Aviti system (Element Biosciences, San Diego, CA).

### Genome assembly, scaffolding, and quality control

We assembled the *Anthemurgus passiflorae* genome using a combination of long-read sequencing and proximity-ligation sequencing. After filtering out adapters from raw HiFi reads using HiFiAdapterFilt v. 0.2.3 (Sim et al. 2022) under default parameters, we performed genome assembly using HiFiASM v. 0.16.1-r375 (Cheng et al. 2021) with the ‘--n-hap 1’ specification. This preliminary contig assembly was assessed for quality and completeness using BlobToolKit v 2.6.1 (Challis et al. 2020) and the Benchmark of Single-Copy Orthologs (BUSCOs; Waterhouse et al. 2017). We used GenomeScope2 to estimate genome size and coverage (Ranallo-Benavidez et al. 2020) and Merqury (Rhie et al. 2020) to assess kmer frequencies. With the HiC data, we scaffolded the contigs into scaffolds using Yet Another Hi-C Scaffolding tool (YAHS; Zhou et al. 2023), which also generates HiC contact maps via the Juicebox tool set. We manually curated the scaffolded assembly in Juicebox (Robinson et al. 2018), then converted the curated scaffold using the Juicebox “juicebox_assembly_converter.py” script. Following conversion of the scaffolded assembly, we re-ran Blobtools and BUSCO to assess assembly quality and identify taxonomic origin of scaffolds. Scaffolds were assigned to their closest taxon using BLAST (Altschul et al. 1997) and Diamond (Buchfink et al. 2015) searches to NCBI nt and UniProt databases, respectively, within BlobToolKit. We removed any scaffolds that were not arthropod nor “no-hit.”

### Mitochondrial genome

Following the MitoHiFi v2 workflow (Uliano-Silva et al. 2023), we identified the mitochondrial genome of *Andrena chekiangensis* (NCBI accession: NC_042768.1) as the closest reference genome. We used Mitos (Bernt et al. 2013) within MitoHiFi to identify and remove the mitochondrial genome from the cleaned, scaffolded whole genome assembly.

### Repeats

In order to characterize repetitive regions within the *P. passiflorae* genome, we implemented RepeatModeler v2.0.4 (Flynn et al. 2020) to identify *de novo* transposable elements. Next, we soft masked *de novo* and known transposable elements from the draft genome assembly using RepeatMasker v4.1.5 (Smit et al. 2024; www.repeatmasker.org/). Both RepeatModeler and RepeatMasker are available within the Dfam consortium software container (https://github.com/Dfam-consortium/TETools/; Storer et al. 2021).

### RNA extraction and sequencing

We sequenced RNA from two tissues to aid in genome annotation. Specifically, total RNA was extracted separately from abdomen and thorax tissues through homogenization of snap frozen tissue in Tri Reagents and then extraction using the Zymo Direct-zol Magbead Total RNA kit (Zymo Research, Irvine, CA) on a Kingfisher Flex 96 system (ThermoFisher, Waltham, MA). From total RNA, a poly(A) cDNA library was generated using the NEB Ultra II RNA Library Prep Kit (New England Biolabs, Ipswich, MA) using oligo(dT) selection. The final RNASeq libraries were sequenced on a PE 150 bp sequencing run on an Element Biosciences Aviti instrument (Element Biosciences, San Diego, CA)

### Genome annotation

Following annotation approaches used in previous bee genome studies (e.g., Sless et al. 2022; Schweizer et al. 2024), we predicted gene annotation using the BRAKER3 workflow (Gabriel et al. 2024). This workflow predicts genes using GeneMark-ETP and AUGUSTUS, using both the *P. passiflorae* thorax and abdomen RNAseq data, in conjunction with the OrthoDB v11 orthologous protein database for Arthropods (Kuznetsov et al. 2022). After annotation with BRAKER3, we performed functional annotation using Interproscan software v5.60-92.0 (Jones et al. 2014; Blum et al. 2020).

### Phylogenetic inference

We constructed a phylogeny of bees and close relatives by extracting Ultraconserved Elements (UCEs) from the iyProPass1 draft genome assembly and a selection of other bee and apoid wasp genomes. To conduct this analysis, we downloaded all publicly available bee genomes from NCBI using the NCBI Datasets command line tools v. 15.6.0 (download date: 23 Feb 2024) and pruned the initial set of genomes to remove duplicate species. We then extracted UCE loci from genomes with Phyluce v. 1.7.3 (Faircloth 2016), following “tutorial 3” from the Phyluce documentation (https://phyluce.readthedocs.io/en/latest/tutorials/tutorial-3.html). UCEs were extracted by aligning the principal v2 UCE probe set for Hymenoptera from Branstetter et al. (2017) to the target genomes and by slicing out 1kb of flanking sequence from either side of each UCE locus. For the probe alignment step, we set both the “min-coverage” and “min-identity” parameters to 80%.

Following UCE extraction, we used a series of Phyluce commands to generate a concatenated UCE matrix for phylogenetic analysis. We ran the phyluce_assembly_match_contigs_to_probes script using the principal version of the probe set and default parameters. Each locus was aligned with MAFFT v.7.475 (Katoh and Standley 2013) and trimmed with Gblocks v0.91b (Talavera and Castresana 2007) using reduced stringency parameters (b1 = 0.5, b2 = 0.5, b3 = 12, b4 = 7). We then filtered the complete set of loci to have 75% taxon completeness and concatenated the alignments into a supermatrix. A phylogenetic tree was inferred using IQ-Tree v. 2.2 (Minh et al. 2020) and a general time reversible model with gamma among site rate variation (GTR+F+G4). To measure support, we conducted 1000 ultrafast bootstrap replicates (UFB; Hoang et al. 2018).

## Results and Discussion

### Assembly

DNA sequencing generated 2,571,591 Sequel reads, of which only 363 (0.014%) were removed with HiFiAdapterFilt. The estimated genome coverage using GenomeScope2 was 150x. The initial HiFiASM assembly consisted of 53 contigs, a L50 of 6, N50 of 19.799 Mb, and a genome size of 251.030 Mb (Figure 2A). Scaffolding with HiC further improved the contiguity of the genome, placing contigs into 49 scaffolds with an N50 of 21.4 Mb. BlobToolKit identified 11 scaffolds of non-Arthropod origin, mapped to the plants *Passiflora lutea, Passiflora edulis, Acacia ligulata, Populus alba*, and *Gossypium hirsutum*, the endosymbiotic bacterium *Wolbachia pipientis*, and the fungus *Meira* sp. Upon removal of these scaffolds, the final genome consisted of 38 scaffolds totaling 249183375 bp, an N50 of 5 Mb, L50 of 21425975 bp, N90 of 11 Mb, and L90 of 13250199 bp (Figure 2B). The assembly has fairly complete Hymenoptera BUSCO, with 97.2% single-copy complete, 0.55% fragmented, 0.30% duplicated, and 2.82% missing (Figure 2B). HiC scaffolding identified 12 chromosomes (Figure 2C).

**Figure 2.**
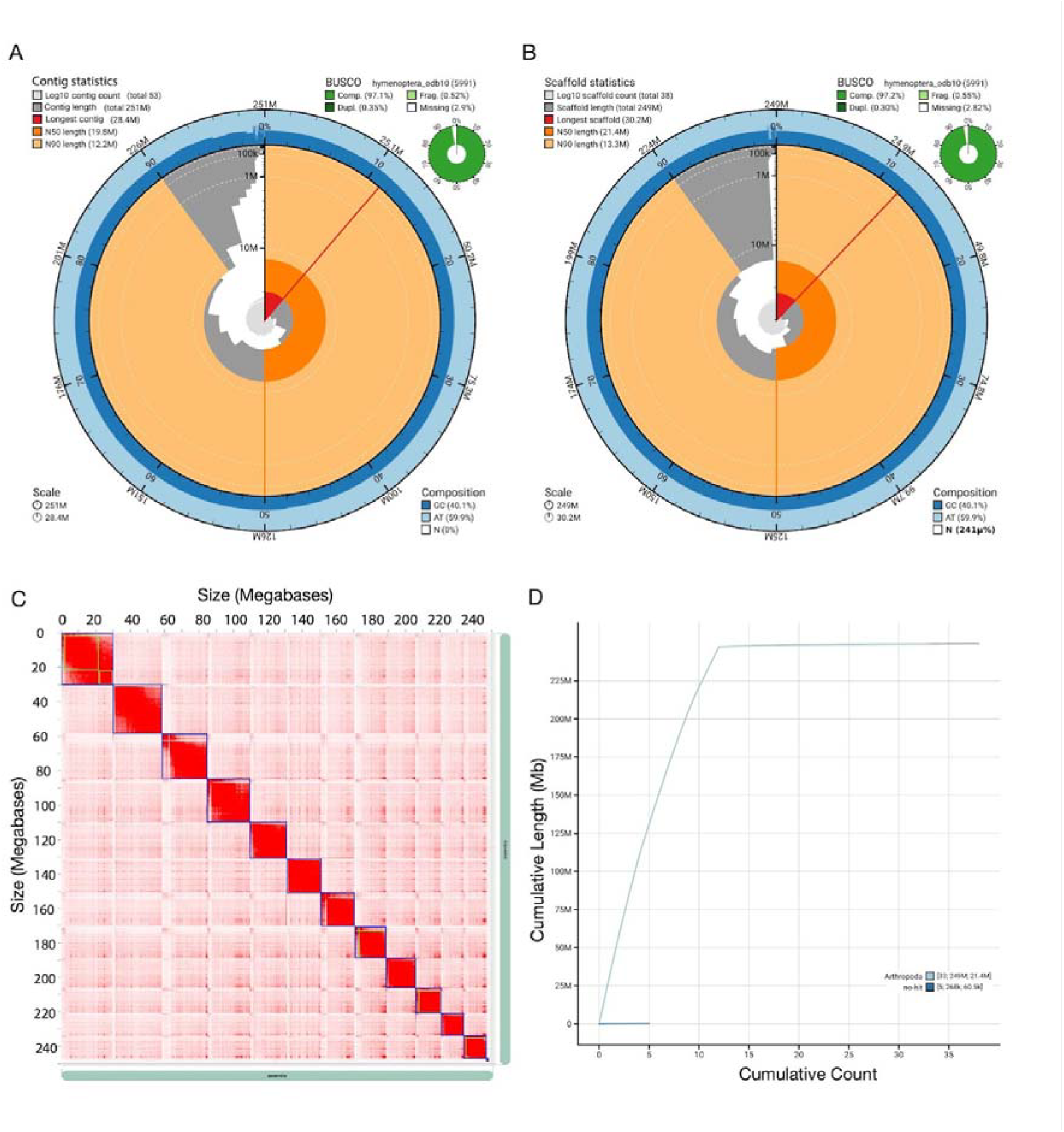
Genome assembly and quality metrics for the *Protandrena passiflorae* genome. A) BlobToolKit snail plot of the initial draft genome contig assembly. The main plot is divided into 1,000 size-ordered bins around the circumference with each bin representing 0.1% of the 251 Mb assembly. The distribution of sequence lengths is shown in dark gray with the plot radius scaled to the longest sequence present in the assembly (31,838,323 bp, shown in red). Orange and pale-orange arcs show the N50 and N90 sequence lengths (17,524,680 and 3,032,719 bp), respectively. The pale gray spiral shows the cumulative sequence count on a log scale with white scale lines showing successive orders of magnitude. The blue and pale-blue area around the outside of the plot shows the distribution of GC, AT, and N percentages in the same bins as the inner plot. A summary of complete, fragmented, duplicated, and missing BUSCO genes in the hymenoptera_odb10 set is shown in the top right. B) Same as A but for the final scaffolded genome. C) HiC contact map of the *P. passiflorae* genome, with 12 chromosomes ordered from largest to smallest, from the top left to bottom right. D) Cumulative length of scaffolded, clean assembly for scaffolds ordered largest to smallest. The two lines indicate taxonomic assignment to Arthropoda (light blue) or no match (dark blue.)

### Repetitive DNA

RepeatModeler and RepeatMasker analyses classified then masked 20.27% of the *P. passiflorae* genome for repetitive elements. Retroelements (1.31%) and DNA transposons (1.63%) were the most common categories of classified repeats, with an additional 15.63% of the genome in unclassified repeats (Table 1).

**Table 1.**
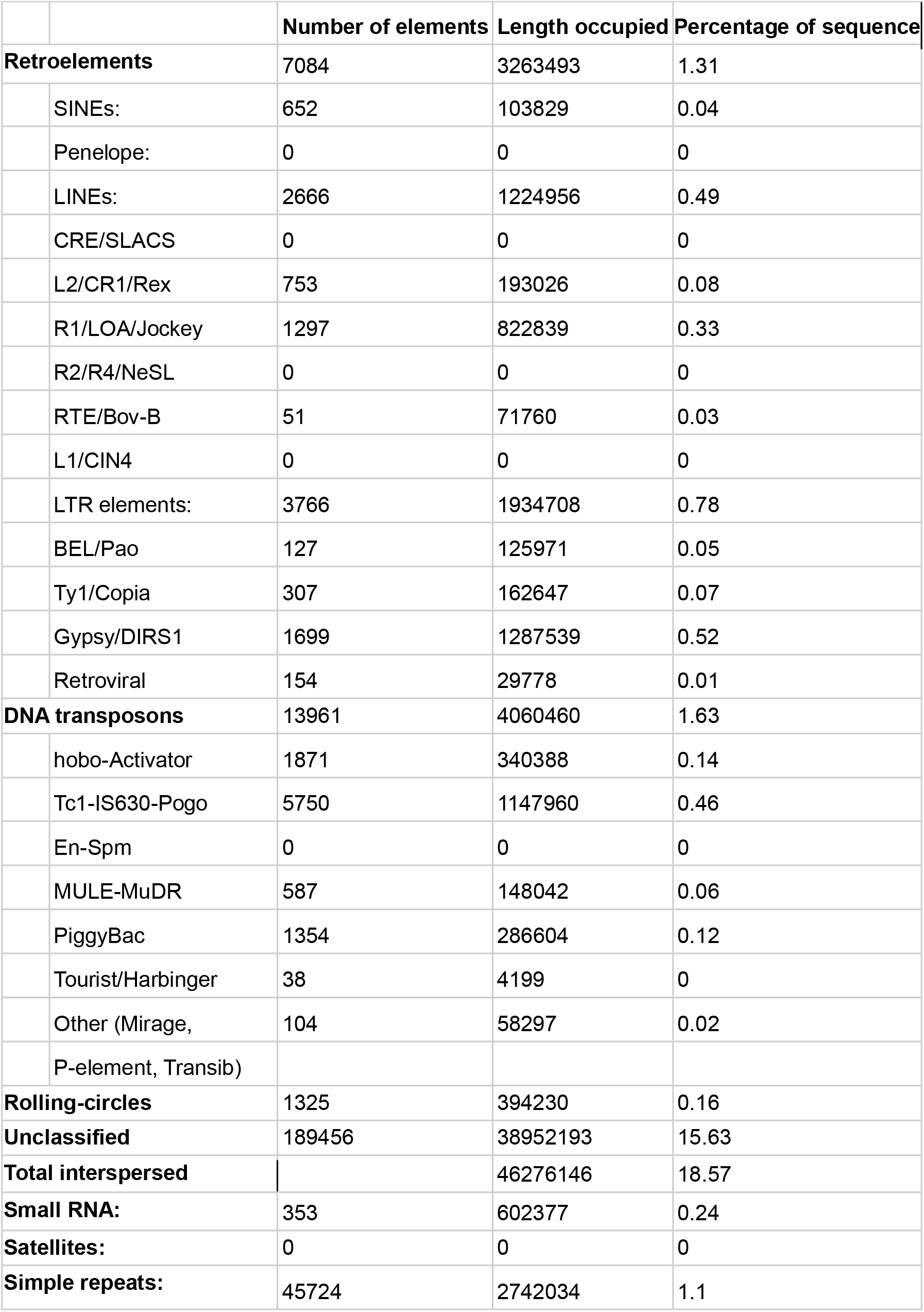
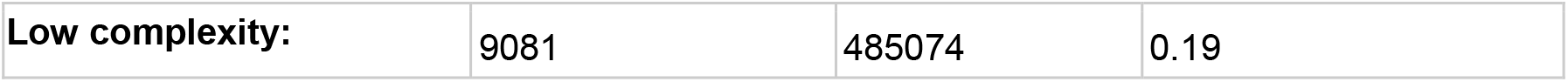
Repetitive elements identified in the *P. passiflorae* genome using RepeatModeler and RepeatMasker.

|  |  | Number of elements | Length occupied | Percentage of sequence |
| --- | --- | --- | --- | --- |
| **Retroelements** |  | 7084 | 3263493 | 1.31 |
|  | SINEs: | 652 | 103829 | 0.04 |
|  | Penelope: | 0 | 0 | 0 |
|  | LINEs: | 2666 | 1224956 | 0.49 |
|  | CRE/SLACS | 0 | 0 | 0 |
|  | L2/CR1/Rex | 753 | 193026 | 0.08 |
|  | R1/LOA/Jockey | 1297 | 822839 | 0.33 |
|  | R2/R4/NeSL | 0 | 0 | 0 |
|  | RTE/Bov-B | 51 | 71760 | 0.03 |
|  | L1/CIN4 | 0 | 0 | 0 |
|  | LTR elements: | 3766 | 1934708 | 0.78 |
|  | BEL/Pao | 127 | 125971 | 0.05 |
|  | Ty1/Copia | 307 | 162647 | 0.07 |
|  | Gypsy/DIRS1 | 1699 | 1287539 | 0.52 |
|  | Retroviral | 154 | 29778 | 0.01 |
| **DNA transposons** |  | 13961 | 4060460 | 1.63 |
|  | hobo-Activator | 1871 | 340388 | 0.14 |
|  | Tc1-IS630-Pogo | 5750 | 1147960 | 0.46 |
|  | En-Spm | 0 | 0 | 0 |
|  | MULE-MuDR | 587 | 148042 | 0.06 |
|  | PiggyBac | 1354 | 286604 | 0.12 |
|  | Tourist/Harbinger | 38 | 4199 | 0 |
|  | Other (Mirage,<br>P-element, Transib) | 104 | 58297 | 0.02 |
| **Rolling-circles** |  | 1325 | 394230 | 0.16 |
| **Unclassified** |  | 189456 | 38952193 | 15.63 |
| **Total interspersed** |  |  | 46276146 | 18.57 |
| **Small RNA:** |  | 353 | 602377 | 0.24 |
| **Satellites:** |  | 0 | 0 | 0 |
| **Simple repeats:** |  | 45724 | 2742034 | 1.1 |

|  |  |  |  |
| --- | --- | --- | --- |
| **Low complexity:** | 9081 | 485074 | 0.19 |

### Gene annotation

The BRAKER3 workflow using orthologous arthropod protein databases and the *P. passiflorae* RNAseq data annotated 12,098 genes and 15,353 total transcripts (Supplemental File: braker.gtf file). The inclusion of the RNAseq data reduced the total number of annotations compared to a BRAKER3 workflow using only the orthologous arthropod protein database (19,553 genes, 21,177 transcripts; results not shown). Interproscan run on the RNAseq and protein workflow was able to assign functional annotation to 10,958 of 12,098 genes and 14,203 of 15,353 transcripts.

### Phylogeny

We recovered an average of 2,455 UCE loci from 131 genomes, including iyProPass1, and generated a concatenated dataset containing 3,566,078 bp of sequence data and 2,526,414 informative sites. We recovered a robust phylogeny of Apoidea that includes a selection of apoid wasp outgroups and all bee families except Stenotritidae (Figure 3). All nodes in the tree except one are highly supported with 100% bootstrap support and relationships among bee families and major lineages generally match recent phylogenomic studies with more balanced taxon sampling (Branstetter et al. 2017, Almeida et al. 2023, Bossert et al. 2021). Within Andrenidae, *Protandrena passiflorae* was correctly placed as sister to the recently published genome of *Perdita meconis* and together they formed a clade sister to *Andrena. Protandrena passiflorae* and *Perdita meconis* both belong to the tribe Panurginae, but are in different tribes, Protandrenini and Perditini, respectively. This divergence represents approximately 54 million years of divergence (Bossert et al. 2021), which explains the long branches connecting the two species. An uncollapsed phylogeny with all 131 genomes is shown in Supplemental file Figure S1. This phylogeny shows that the taxonomic sampling of Andrenidae for full reference genomes remains incomplete and highly biased toward the single genus *Andrena*. By adding a new tribe to the tree, we fill in an important gap in sampling.

**Figure 3.**
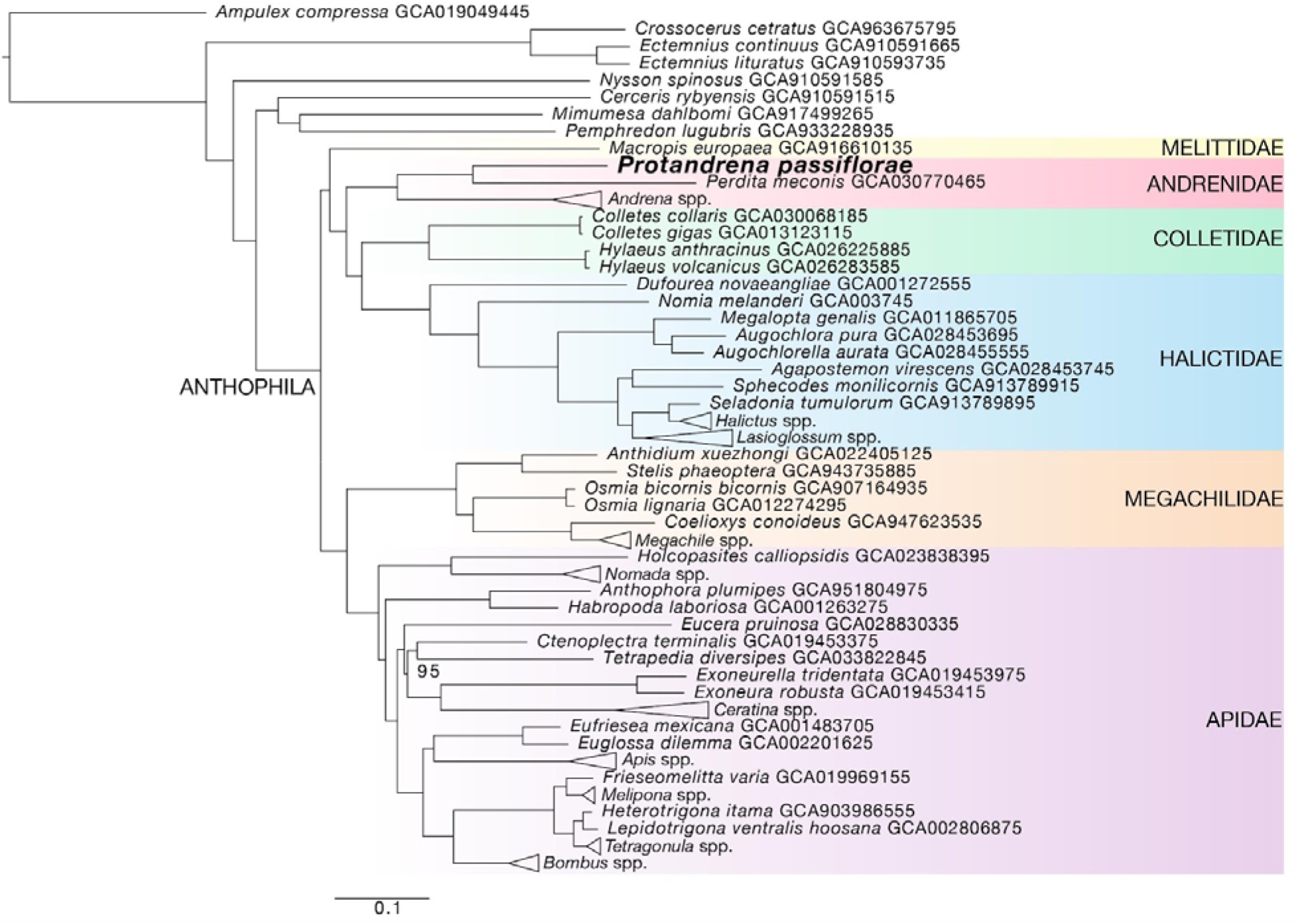
Phylogenetic tree based on the analysis 2,487 ultraconserved element loci extracted from 131 genomes, including the target species *Protandrena passiflorae*. The tree was inferred with IQ-Tree and support was estimated using ultrafast bootstrap replicates. Bootstrap values are only shown if less than 100%.

## Summary

### Data availability

This Whole Genome Shotgun project has been deposited at DDBJ/ENA/GenBank under the accession JAZGQT000000000. The version described in this paper is version JAZGQT010000000. Raw reads from HiFi, HiC, and RNAseq have been deposited in the NCBI Short Read Archive under BioProject PRJNA1072036.

## Acknowledgments

The genome assembly was generated as part of the USDA-ARS Beenome100 Initiative (https://www.beenome100.org/). The authors thank the members of the USDA-ARS Beenome100 and Ag100Pest Team for sequencing and analysis support. All opinions expressed in this paper are the authors’ and do not necessarily reflect the policies and views of USDA. Mention of trade names or commercial products in this publication is solely for the purpose of providing specific information and does not imply recommendation or endorsement by the U.S. Government. USDA is an equal opportunity provider and employer.

