## Supplementary figures and images for "Chromosome-level genome assembly of *Protandrena* (*Anthemurgus*) *passiflorae* (Hymenoptera: Andrenidae), a host-plant specialist bee"

### Supplemental file Figure S1

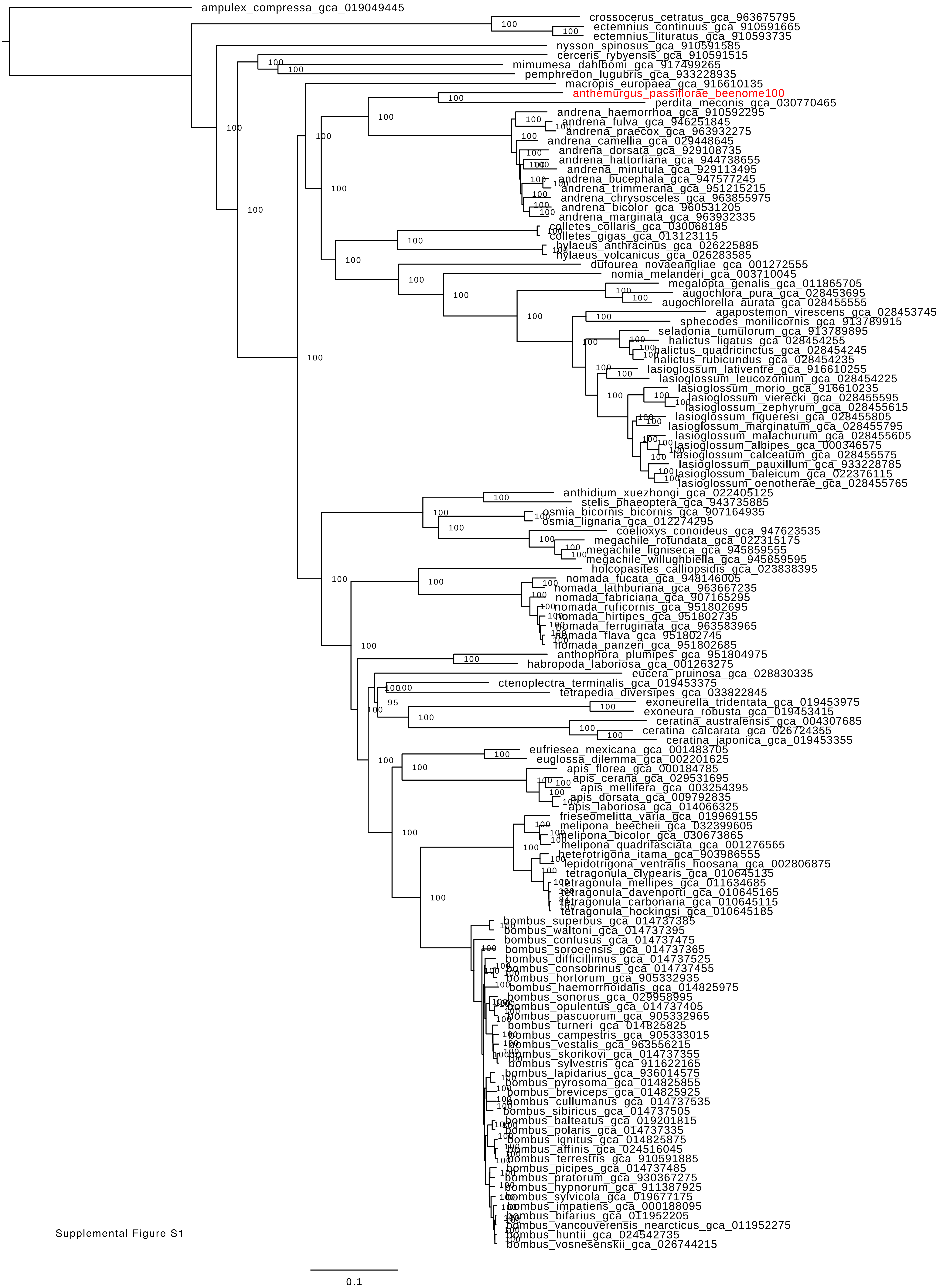


Supplemental Figure S1
